# Embodied mechanical sensing through self-generated motion in climbing plants

**DOI:** 10.64898/2026.03.16.712031

**Authors:** Amir Ohad, Amir Porat, Yasmine Meroz

## Abstract

Embodied mechanical intelligence and morphological computation enable adaptive behavior by shifting computation from centralized control to body mechanics. Yet how dynamical material structures encode environmental information remains poorly understood. Here, we address this question using climbing plants as a minimal decentralized system. Combining force measurements with physical modeling, we show that circumnutating shoots generate predictable forces against candidate supports, and that the interaction is equivalent to a rotating cantilever governed by torque balance. Force amplitude is determined by stem stiffness and geometry, while the timescale is set by circumnutation rate. Twining occurs only after reaching a critical torque threshold, corresponding to a deformation that reflects support stability and geometric reach. Motorized perturbations further show that twining time scales inversely with effective circumnutation rate. Together, these results show how self-generated motion extracts actionable information about the environment, while the tapered geometry and spatially varying mechanics of the shoot encode the decision-making threshold for attachment.

## Introduction

Embodied mechanical intelligence and morphological computation offer a powerful alternative to centralized control, offloading computational complexity from an organism’s or robot’s “brain” to its physical body (*1–4*). By leveraging intrinsic material properties, geometry, and environmental interactions, such systems achieve adaptive behavior, energy efficiency, and robustness without the need for complex control algorithms. While these principles are foundational to the development of soft robotics and intelligent matter, identifying the underlying physical laws that govern decentralized decision-making remains a substantial challenge. Existing biological examples of active sensing, such as the whisking behavior of mammals (*5–7*), are deeply integrated with complex neural architectures and sophisticated anatomy, making it difficult to decouple the role of pure mechanics from neural processing. To isolate the fundamental principles of embodied sensing, a simpler, fully decentralized model system is required.

Climbing plants provide such a system. In the race for access to light, these plants prioritize rapid vertical growth over radial thickening, sacrificing mechanical stability in favor of speed (*8, 9*). Lacking the capacity to support their own weight, they must attach to external objects, tying their survival to the ability to locate and select structurally sound supports. Because they lack a nervous system or central controller, their behavior is entirely emergent from the coupling of their growth-driven mechanics and environmental feedback (*8, 10*). Furthermore, their anatomy can be approximated as a slender, active material rod, offering a minimal biological platform to investigate how mechanical structure alone can encode functional responses. It is well established that climbing plants use circumnutations (CN), self-generated periodic oscillatory movements, to explore their surroundings (*11–13*). However, the physical nature of the information extracted during these oscillations has remained largely unexplored. We hypothesize that circumnutations function as a mode of active mechanical sensing: by generating predictable periodic loading against external objects, these oscillations enable the plant to probe a support’s mechanical stability and its own capacity to achieve a firm grasp. This mechanism allows the system to distinguish between a rigid, load-bearing support and an unstable one through purely mechanical feedback, triggering twining only when specific physical criteria are met.

Though the biomechanics of twining are well-documented (*8, 14–18*), the mechanical interactions that precede stable attachment, where the “decision” to twine is made, remain largely unexplored (*15, 19*). We address this by focusing on the common bean (*Phaseolus vulgaris*) as a model system and modeling the circumnutating shoots as rotating Cosserat rods. By combining high-resolution force measurements with morpho-elastic simulations, we show that the interaction reduces to a minimal physical model: a rotating cantilever governed by torque balance, where force amplitude is determined by shoot geometry and stiffness. The conceptual flow of the study is summarized in Supplementary Fig. S1.

Our results reveal that the plant’s tapered geometry and elastic gradient are functionally tuned for sensing: twining initiation depends jointly on support stability and contact geometry, requiring a minimal tip overshoot for a successful grasp and occurring only after a critical bending moment threshold is reached. Together, these results provide direct evidence for active mechanical sensing in plants. Understanding these principles demonstrates how decentralized systems utilize active mechanics to achieve robust, autonomous decision-making, providing a blueprint for engineered soft robots and adaptive materials that can sense and act through their inherent mechanical structure.

## Results

### Circumnutations exert predictable forces on candidate supports

We measured the interaction force between a circumnutating bean stem and a support (n=164) using a previously developed physical pendulum force sensor (*20*). In this setup, a bean plant is placed so that during its circumnutation movement it will reach the pendulum rod and push it. Fig. 1A shows a schematic of the experimental setup, where the total extent of the shoot is divided into the lever length*ℓ*_lev_, from the base of the plant to the point of contact, and the overshoot length *ℓ*_o_, from the point of contact to the tip. The variable *s* follows the centerline, with *s*=0 at the tip, and we define the point *s*_*c*_ = *ℓ*_o_ at the point of contact. The force is calculated from the rod deflection angle *α* as a function of time. An example of a circumnutating bean deflecting a pendulum rod is shown in Fig. S2 and Movie S1. Examples of measured force trajectories are shown in Fig. 1B, truncated when the plant either twines or slips past the rod. All force trajectories can be found in Fig. S3. Force trajectories exhibit a large range of forces and durations, with force amplitudes ranging between 0.1 mN and 2.7 mN.

**Figure 1:**
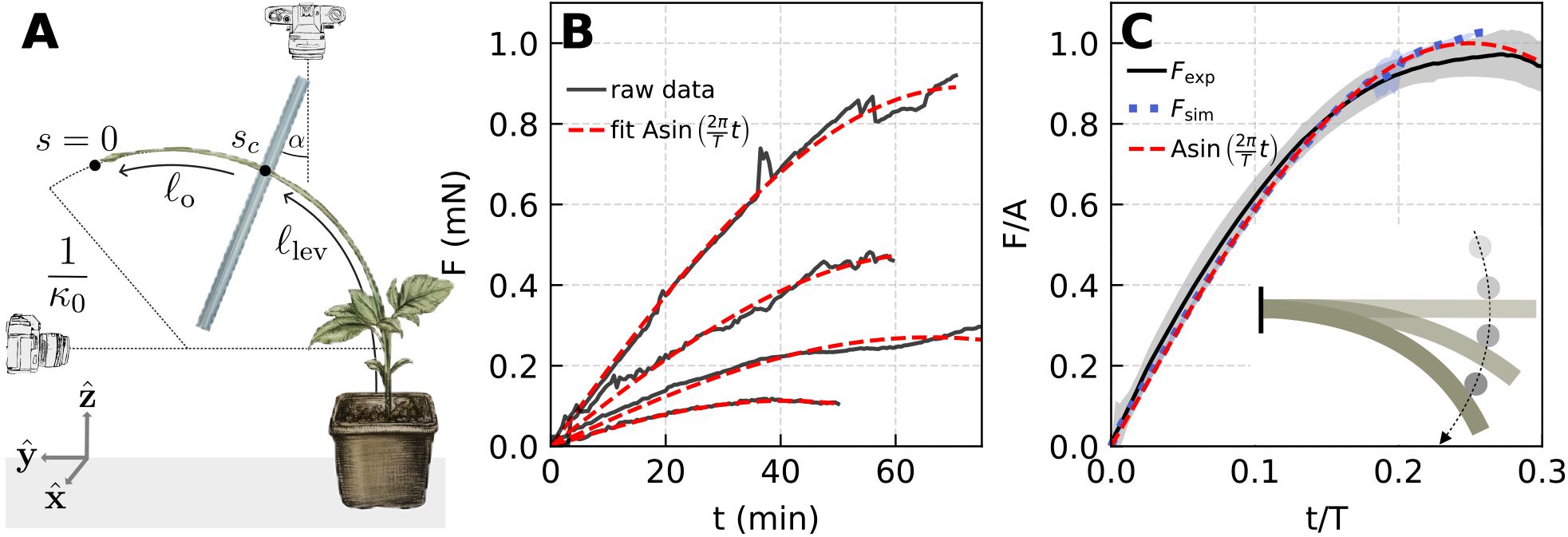
Force measurements of a circumnutating bean stem interacting with a pendulum support. **A**. Setup schematic (after Darwin’s illustrations). The circumnutating stem deflects the pendulum rod an angle *α*, which we translate to the pushing force*F*. We define the variable*s* following the stem centerline, with *s*=0 at the tip. The point of contact with the rod is *s* = *s*_*c*_. We define the distance from the tip to the point of contact as the overshoot length *ℓ*_o_, and the distance from the point of contact to the base as the lever arm length *ℓ*_lev_. The intrinsic curvature of the stem *κ*_0_. Imaging is performed using side and top views. **B**. Examples of measured raw force trajectories (solid black curves). Smoothed trajectories are fit to a sine function *A* sin (2*πt* / *T*) (dashed red curves). **C**. Normalized force trajectories. Force and time of measured trajectories (n=164) are normalized by the fitted amplitude *A* and period *T*, and averaged (solid black curve). The shaded region shows the standard deviation (SD). Simulations are normalized in the same manner (see Fig. S7). Simulation variability is computed as the SD of normalized forces for parameter ranges of the Young’s modulus *E* ∈ [10,100] MPa, friction coefficient *µ* ∈ [0,1], and *ℓ*_o_ / *L* ∈ [0.1,0.54] (dotted blue curve and shaded region). A sine function is shown for reference (dashed red curve). (inset) The interaction is well described by a rotating cantilever with a fixed load (Fig. S8), or equivalently, in the plant frame, a cantilever (green) subjected to a rotating load (grey), shown schematically.

Forces evolve following a characteristic sinusoidal trajectory, as evident from the good fit to a sine function for the example trajectories in Fig. 1B (distribution of fit scores in Fig. S4). Normalizing trajectories by the fit sine amplitude and period (see Methods) collapses them onto a common trajectory (Fig. 1C). The consistent periodic force generation observed in our experiments is indicative of active mechanical sensing, in which self-generated, controlled motion enables the extraction of information that cannot be obtained from passive interaction alone. Examples of such active sensing strategies include whisking in cats and rodents (*5–7*); insect antennae (*21*); flagellar beating in microorganisms (*22, 23*).

### Mechanically informed minimal model recapitulates measured sinusoidal force generation

To identify the physical mechanism underlying this predictable force generation, we developed a minimal, experimentally informed model describing the mechanical interaction observed in the experimental setup. Slender stems are represented as Cosserat rods (*24*), which are 1D elastic continuous elements able to undergo all modes of deformation - bending, twisting, stretching and shearing - and reconfigure in 3D space. We define a variable *s* that follows the stem centerline, with *s* =0 at the tip, and we mark the point of contact with rod *s*_*c*_. In our minimal model we assume the shoot is of constant length *L*, with a uniform intrinsic curvature *κ*_0_ (Fig. 1A), and the Young’s modulus *E*, shoot radius *R* are constant along the shoot.

To inform this model, after each of the force measurements described in the preceding section, we performed a quantitative characterization of the geometric and mechanical properties of each plant stem (details in Methods). We measured the radius *R* and Young’s modulus *E* (Fig. 2) of consecutive 5-cm segments from the apical tip to 50 cm below. The radius of the stem increases linearly with distance from the tip, consistent with other measured plant shoots (*25–31*). The average value at the point of contact across all shoots is 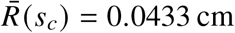 (Fig. 2B). The Young’s modulus increases parabolically away from the tip, consistent with prior measurements (*27, 28, 30, 32–35*), reflecting changes in shoot composition as a result of tissue maturation further from the tip. The average Young’s modulus at the point of contact is *Ē*(*s*_*c*_)=2.6 MPa (Fig. 2C). To further inform the model, we characterized both the free circumnutation dynamics prior to contact and the contact geometry with the support. The circumnutating shoots exhibited a characteristic period of approximately 100 min, while the spatial extent of the circumnutation dynamics was characterized by the maximal intrinsic stem curvature, *κ*_max_. The relatively narrow distribution of curvature values (Fig.S5D), together with representative stem geometries (Fig.S8), is consistent with the assumption of approximately symmetric circumnutation adopted in the minimal model. A video of a circumnutating stem can be found in Movie S2. We further measured the overshoot length *ℓ*_o_ and lever arm length *ℓ*_lev_ at the onset of contact. Average values of all measured parameters are summarized in Table 1, with their corresponding distributions shown in Fig. S5.

**Table 1:** Table of average measured values of parameters, and simulation values. The simulation parameters were chosen to be of the same order of magnitude as the measured values, though not identical, in order to reduce computation time, as small variations can substantially affect simulation speed. The normalized simulation trajectories remain invariant over broad ranges of Young’s modulus, friction coefficient, and overshoot length (see Fig. 1C and Fig. S1), demonstrating that the characteristic sinusoidal dynamics are robust to parameter variation despite the use of representative simulation values. Radius *R* and Young’s modulus *E* are extracted values at the point of contact with the support (Fig. 2). Other experimental values taken from distributions in Fig. S5. Intrinsic curvature *κ*_0_ is taken from the maximal measured curvature *κ*_max_. Errors are the standard deviations.

| Experimental measurements | Representative simulation parameters |
| --- | --- |
| $\bar{R}(s_c) = 0.043 \pm 0.001 \text{ cm}$ | $R = 0.05 \text{ cm}$ |
| $\bar{E}(s_c) = 2.6 \pm 3.3 \text{ MPa}$ | $E = 10, 50, 100 \text{ MPa}$ |
| $\bar{T}_{\text{cn}} = 98 \pm 11 \text{ min}$ | $T_{\text{cn}} = 100 \text{ min}$ |
| $\bar{\ell}_o = 2.1 \pm 1.7 \text{ cm}$ | $\ell_o = 0.55 - 2.7 \text{ cm}$ |
| $\bar{\ell}_{\text{lev}} = 42 \pm 8 \text{ cm}$ | $\ell_{\text{lev}} = 2.3 - 4.45 \text{ cm}$ |
| $\bar{\kappa}_{\max} = 0.16 \pm 0.05 \text{ cm}^{-1}$ | $\kappa_0 = 0.4 \text{ cm}^{-1}$ |

**Figure 2:**
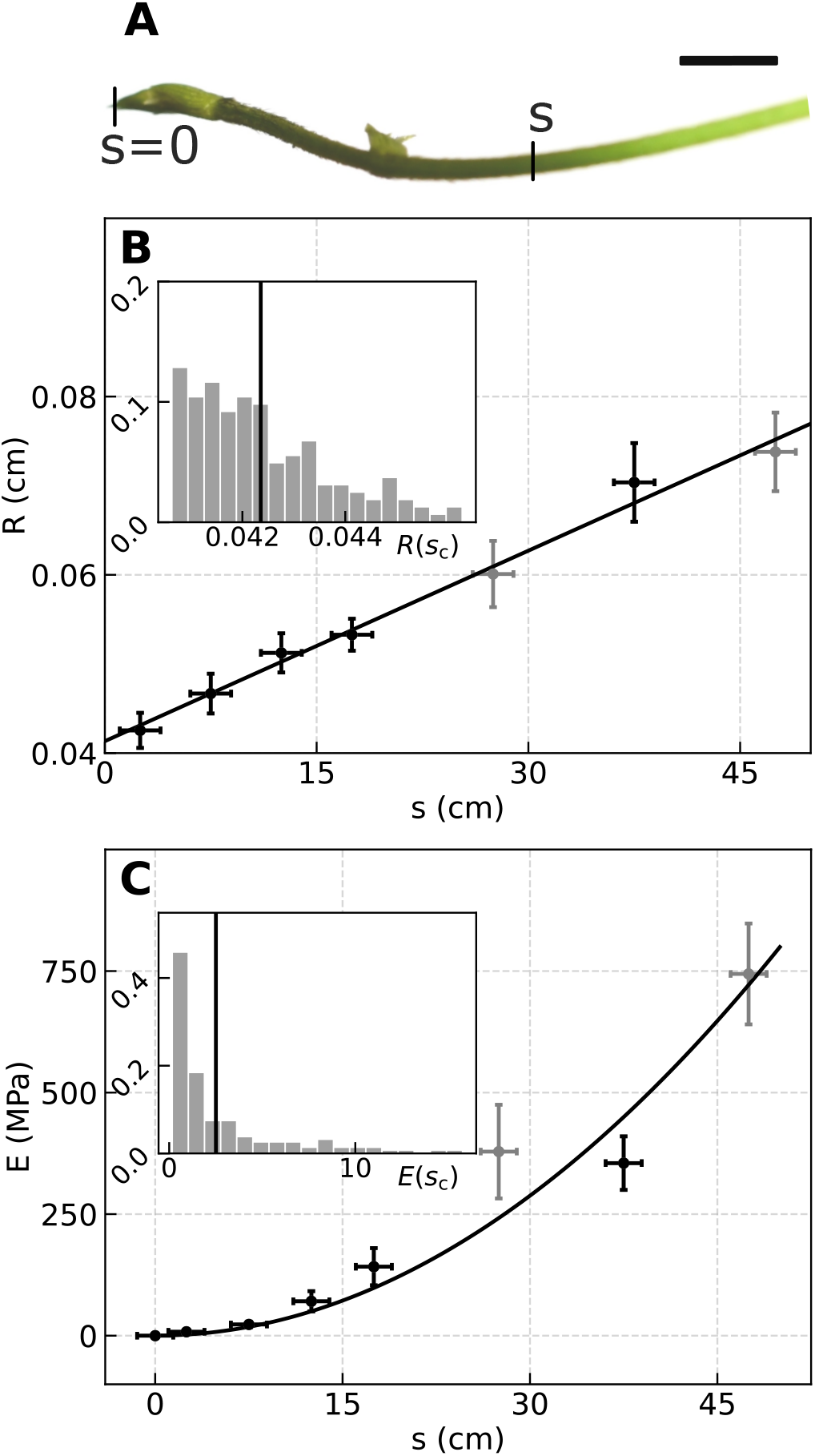
Morphological and mechanical characterization of the plant stem. **A.** Close-up image of a representative bean stem. The coordinate *s* follows the centerline, with *s* = 0 at the tip (Fig. 1A). Scale bar: 1 cm. **B**. Average stem radius *R* (*s*) along the centerline (*n* = 42, 51, 53, 51, 2, 8, 2). The linear fit was performed excluding the two light gray symbols (n=2), which are shown for completeness only. A linear fit yields *R* (*s*) = 0.0008 *s* + 0.0406 cm, *R*^2^ = 0.99. Error bars show the standard error across samples. (inset) Distribution of radius values at the point of contact with the rod *R*(*s* = *sc*) (n=106). Vertical line indicates mean value, 0.0429 ± 0.0012 cm. **C**. Average value of Young’s modulus *E* (*s*) along the centerline (same n values as in **B**). A parabolic fit yields *E* (*s*) = 0.30 *s*^2^ + 0.20 MPa, *R*^2^ = 0.91. Distribution of Young’s modulus values at the point of contact *E* (*s* = *s*_*c*_) (n=106), with a mean value 2.6 ± 1.6 MPa.

Building on our physical model, informed by experimental values in Table 1, we ran simulations (see Fig. S7) using *Elastica* (*24,36*), a numerical solver for Cosserat rods (see Methods). The physical parameters were chosen to be of the same order of magnitude as the measured values, though not identical, in order to reduce computational time without loss of generality, as small variations can substantially affect simulation speed.

A separation of timescales between slow growth-driven circumnutations and fast mechanical relaxation allowed us to decouple the two processes in a quasi-static manner (*37–39*). We emulated circumnutations (driven by a rotating differential growth gradient in the cross-section) by fixing the base of the stem, and rotating its curvature with the circumnutation period *T*_cn_ (without rotating the material frame). A rod of infinite stiffness, representing the support, was placed at different positions relative to the stem reach, yielding different overshoot and lever lengths *ℓ*_o_ and *ℓ*_lev_. When the stem came into contact with the rod, the simulation provided the evolution of forces applied by the stem rotating onto the rod. A video of a typical simulation can be found in Movie S3, and a graphical rendering in Fig. S6, with force simulation trajectories shown in Fig. S7.

In all, the minimal model neglects growth and spatial variations of *R,E*, and *κ*_0_, yet captures the observed sinusoidal force dynamics (Fig. 1C). Importantly, the normalized simulation trajectories remain invariant over broad ranges of Young’s modulus, friction coefficient, and overshoot length (see Fig. 1C and Fig. S1), demonstrating that the characteristic sinusoidal dynamics are robust to parameter variation despite the use of representative simulation values. Building on this agreement, we seek a minimal analytical description of the interaction. Because both the measured circumnutation motion and the dominant force component lie in the horizontal 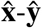 plane, we can further simplify this interaction by considering its projection onto this plane. The interaction can therefore be understood as a rotating cantilever with a fixed load. Equivalently, in the frame of reference of the plant, to a cantilever subjected to a rotating load (detailed in Fig. S8, and inset Fig. 1C).

For a standard cantilever with a load that translates at constant speed *v*, the force grows linearly as *F*(*t*)=*k v t*, where *k* is the beam’s spring constant. In contrast, when the load rotates with angular frequency Ω, only the component perpendicular to the beam induces deflection. The resulting force therefore follows:

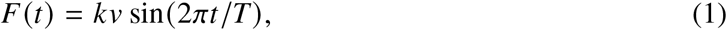

in line with the observed force trajectories.

Having validated that the minimal model reproduces the characteristic force dynamics (Fig. 1C), we built on the analogy to a cantilever and proceeded to use it as an *in silico* laboratory to systematically vary the governing mechanical parameters. This allows us to test the robustness of the proposed physical mechanism and identify the variables responsible for the observed variability in force trajectories.

### Torque balance identifies variables governing variability in measured force trajectories

Following the cantilever analogy, together with our quasi-static assumption, equilibrium requires a balance between the external moment generated by the circumnutating plant, equal to a force acting at a lever arm *M*_ext_ =*Fℓ*_lev_, and the internal restoring or bending moment resulting from the stem’s resistance to bending curvature, defined as *M*_int_ = *E I*(*κ*−*κ*_0_), where *κ*_0_ and *κ* are the intrinsic and actual curvatures of the stem, respectively (Fig. S9 shows a schematic). *B* = *E I* is the flexural rigidity, where *E* is Young’s modulus, and 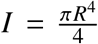 is the area moment of inertia of the stem, approximated as a rod with radius *R*. Thus, torque balance *M*_ext_ =*M*_int_, relates the applied force directly to the mechanical properties of the shoot and its deformation:

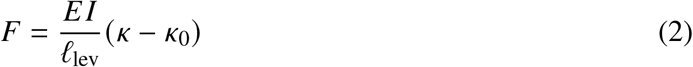

This relation predicts that the applied force increases linearly with higher values of Young’s modulus *E*, or inversely with lever arm *ℓ*_lev_. We tested these predictions both *in silico* and experimentally. We carried out simulations for a range of values of the Young’s modulus, *E* =10,50,100 MPa, and for each we placed the support at different distances from the plant base, yielding a range of lever length ratios *ℓ*_lev_/*L* ∈ [0.4,0.9], where *L* is the total length of the stem.

To evaluate the dependence of force on Young’s modulus, we fixed *ℓ*_lev_/*L* = 0.85 and varied *E*. Plotting the maximal force of each trajectory, *F*_max_, against the corresponding Young’s modulus *E* reveals a clear linear dependence (Fig. 3A), consistent with Eq. 2. The full force trajectories are shown in the inset. Experimentally, we binned force trajectories by the Young’s modulus at the contact point *E* (*s*_*c*_)(n = 164) and averaged the trajectories within each bin (representative trajectories shown in Fig. 3B). Plotting the maximal value of the averaged force, *F*_max_, against the Young’s modulus is consistent with the predicted linear scaling. (Fig. 3B). This pattern corresponds to the intuitive understanding that a stiffer stem deforms less and therefore pushes the support with greater force. Slight differences between predicted and measured values of the forces may be accounted for by recalling that simulations assume a minimal model, e.g., assuming a constant *E* and *R* along the stem. Unlike the simulations, the experimental parameters are coupled, since the local Young’s modulus and stem radius depend on the position of the contact point, corresponding to the overshoot or lever arm length (Fig. 2). Thus, simulations isolate individual parameter effects, whereas experiments reflect their combined influence. Furthermore, we note that simulations provide the total force, while the experimental measurement provides a 2D projection of the force on the 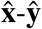plane. Since this plane is parallel to the circumnutation movement, forces in the 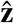 direction are negligible. In Fig. S10 we compare the total simulated force *F* to the projection *F*_*xy*_, finding that the difference between the two increases with friction, and particularly as the support comes into contact farther from the tip (i.e a larger *ℓ*_o_/*ℓ*_lev_ ratio), since the stem is more vertical, and the 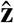 component increases.

**Figure 3:**
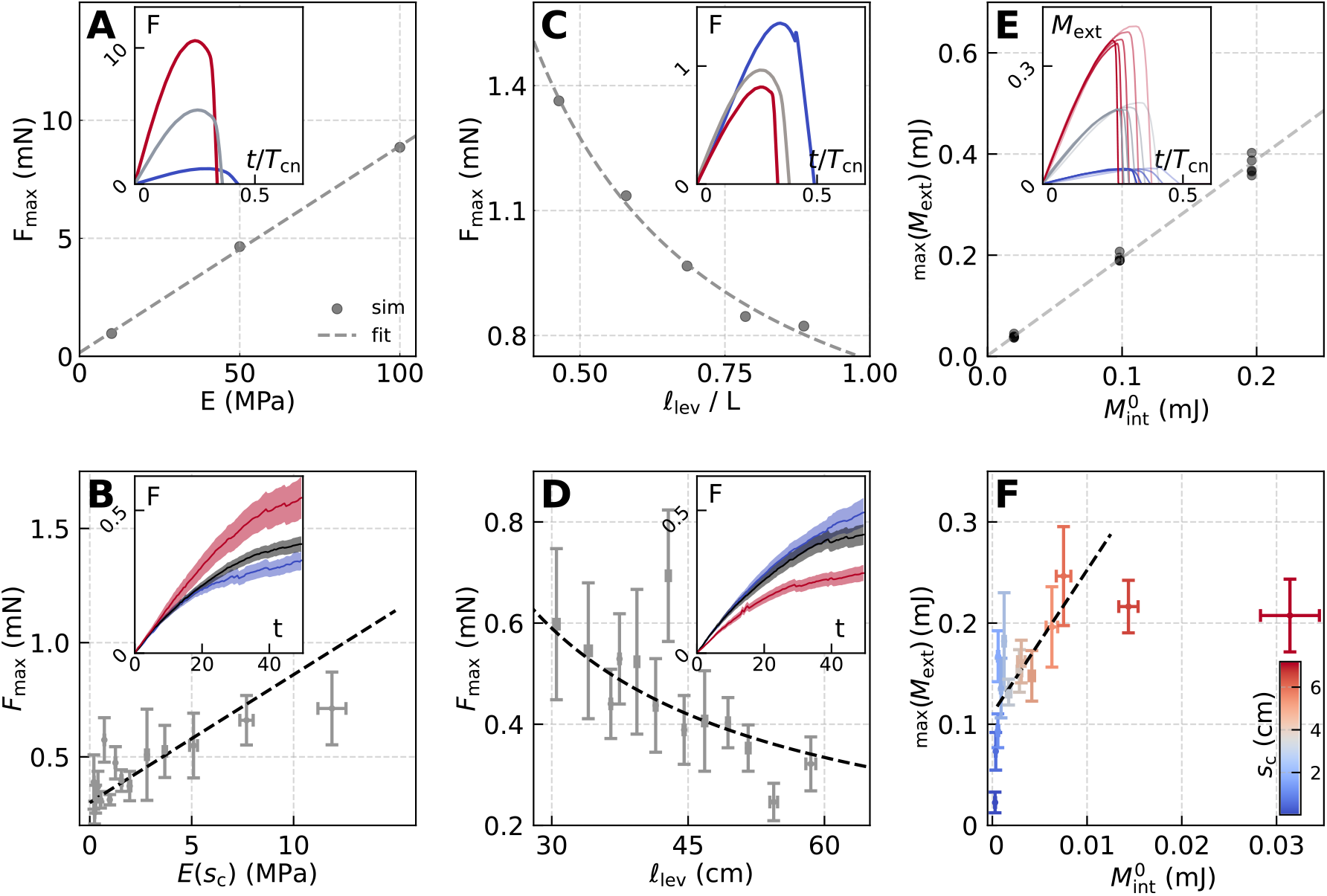
Simulations identify physical variables governing variability in measured force trajectories. **A.** Maximal force for each trajectory *F*_max_ vs Young’s modulus *E*. A linear fit yields *F*_max_ = 0.08 *E* + 0.15 mN, with *R*^2^ = 0.99. All simulation force trajectories (including friction) can be found in Fig. S7. (inset) Simulated forces *F* (without friction) as a function of time *t* normalized by the simulated CN period *T*_cn_, for organs with *ℓ*_o_ / *L* = 0.15. Colors denote the Young’s modulus for values *E* ∈ 10, 50, 100 MPa (blue, grey, red). **B**. Mean maximal force *F*_max_ (obtained as the fitted sine amplitude) binned uniformly (*n* = 11 per bin) according to the Young’s modulus at the contact position *E* (*s*_*c*_). Error bars represent one SD. A linear fit yields *F*_max_ = 0.058 *E* + 0.298 mN, with *R*^2^ = 0.4 and 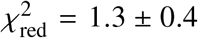 (inset) Experimental force trajectories grouped according to *E* (*s*_*c*_) ∈ (0 − 1, 1 − 4, 4 − 17) MPa (*n* = 49, 82, 33). Shaded regions represent SD around the mean trajectory. **C**. Maximal force *F*_max_ as a function of *ℓ*_lev_, fitted to an inverse function of *ℓ*_lev_/*L* (dashed line) yielding *F*_max_ = 0.35*L*/*ℓ*_lev_ − 1.34 mN with *R*^2^ = 0.99. (inset) Simulated forces as a function of normalized time for *E* = 10 MPa and *ℓ*_lev_/*L* ∈ 0.5, 0.7, 0.9 (blue, grey, red). **D**. *F*_max_ binned uniformly (*n* = 11) by lever length *ℓ*_lev_ ∈ (30, 60) cm. Fitting to an inverse function of *ℓ*_lev_ yields 15.32 *L*/*ℓ*_lev_ + 0.07 mN, with *R*^2^ = 0.5 and 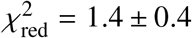. (inset) Experimental force trajectories for *ℓ*_lev_ ∈ (28, 38], (38, 52], (52, 62] cm (*n* = 49, 82, 33, blue, grey, red). **E**. The maximal external torque *M*_ext_ = max(*Fℓ*_lev_) in the simulations scales linearly with the internal bending moment *M*_int_ = *E I κ*_0_. The dashed line is the linear fit: *M*_ext_ = 1.937*M*_int_ + 0.001 mJ, with *R*^2^ = 0.99 and 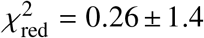. The data points represent simulations with *E* ∈ 10, 50, 100 MPa. (inset) External torque *M*_ext_ as a function of time for the range of *E* values (colors as in A) and *ℓ*_lev_ ∈ [28, 62] cm (opacity increases with *ℓ*_lev_). **F**. Estimated maximal external torque max(*M*_ext_) = max(*Fℓ*_lev_) as a function of the internal torque assuming 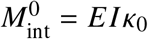, for trajectories grouped by *s*_*c*_ (colorbar, *n* = 12). Linear fit yields *M*_ext_ = 15.66 *M*_int_ + 0.11 mJ, with *R*^2^ = 0.5 and 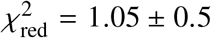. The assumption *κ* = 0 does not hold for small *s*_*c*_, and saturation at large *s*_*c*_ may reflect a maximum torque the plant can generate.

To probe the dependence on the length of the lever arm *ℓ*_lev_, we fixed the Young’s modulus *E* and varied *ℓ*_lev_. Plotting *F*_max_ of each trajectory against the corresponding *ℓ*_lev_ value clearly shows that force *F* depends inversely on lever distance (Fig. 3C), consistent with the torque balance in Eq. 2. Experimentally, we restricted trajectories to *E* (*s*_*c*_) ∈ (0,17)MPa, binned them by *ℓ*_lev_, and averaged within each bin. The maximal averaged force *F*_max_ plotted against *ℓ*_lev_ recovers the predicted inverse dependence. Representative force trajectories are shown in the inset.

Having corroborated the predicted scaling of*F* with *E* and *ℓ*_lev_, we tested the validity of the torque balance in Eq. 2, comparing the external moment *M*_ext_ = *Fℓ*_lev_ to the internal bending moment *M*_int_ = *E I*(*κ*−*κ*_0_).

Since in experiments we do not have access to the actual bending curvature*κ*, we approximate the relation for the case when a stem is being straightened due to the interaction with the support (*κ*=0), as reflected in Fig. S6. As a basis for comparison with experimental results, we plotted the maximal value of external torque max (*M*_ext_), for each of the simulations in Fig. 3B, D, against the respective internal restoring torque assuming 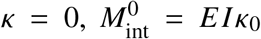 (Fig. 3E), recovering a linear relation consistent with the torque balance in Eq. 2. As expected, simulations with different contact positions, i.e., different values of *ℓ*_lev_, collapsed to similar values, according to their Young’s modulus value *E*. This is further elucidated in moment trajectories *M*_ext_(*t*) (inset), which collapse onto similar values, modulated by the Young’s modulus.

To corroborate these results experimentally, we grouped trajectories according to the contact point position *s*_*c*_, since it provides a direct correspondence to the local parameters 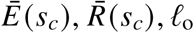, and *ℓ*_lev_. For each experiment, we used the measured values specific to that plant, namely *ℓ*_lev_ and *F*to compute *M*_ext_, and *E*(*s*_*c*_), *R*(*s*_*c*_), and *κ*_0_ = *κ*_max_ to compute 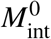. The resulting experimental torque relation is shown in Fig. 3F, revealing a linear dependence over an intermediate range of contact positions *s*_*c*_. For large values of *s*_*c*_, we observe a saturation of *M*_ext_, which may reflect a maximum limit on growth-driven force production. Furthermore, larger *s*_*c*_ correspond to smaller *ℓ*_lev_, where the neglected 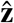 component of the measured force *F* becomes non-negligible (Fig. S10). For small *s*_*c*_, the rapid increase in *M*_ext_ may arise because the simplifying assumption *κ*=0 is not valid for short overshoots *ℓ*_o_, which only deflect at the tip.

In all, these experiments and simulations support our proposition that a circumnutating shoot pushing at a candidate support is analogous to a clamped cantilever with a rotating load, where torque balance is retained. In particular, the amplitudes of the measured force trajectories are mainly modulated by the point of contact along the stem, through its effect on local stiffness *E* and lever arm *ℓ*_lev_.

### Twining is initiated according to stability of support and optimal grasp positioning

Having elucidated the mechanical interaction between the circumnutating stem and the candidate support, we now turn to identify the physical signal that sets twining initiation. To this end, we reexamined the data from the experiments described in the previous section at the level of individual contact events, classifying each according to their outcome: twining or slipping, as described in the Methods section (see twining initiation example in Movie S1.Our first step was to validate the underlying hypothesis that climbing plants initiate twining on a newly encountered support based on the evaluated mechanical stability. The structural stability of a support is reflected in the force it exerts in response to an external load, e.g., a circumnutating plant. Therefore, for each experiment we calculated the resistance of the pendulum rod (the support candidate in our experimental setup). Based on torque equilibrium (*20*), the resistance follows 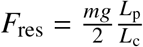, where *m* and *L*_p_ are the mass and length of the pendulum respectively, g is the gravitational acceleration, and *L*_c_ is the distance of the point of contact with the plant from the top of the pendulum, which we gathered for each experiment, as detailed in (*20*). Because the support resistance depends on the pendulum mass, the measurements performed with different pendulum masses (Methods) naturally span a range of resistance values. For further robustness and to increase the range of resistance values, we carried out additional sets of similar experiments, using rods with different masses (0.08 gr and a fixed support (*F*_res_ → ∞)). Fig. 4A shows that the fraction of twining events increases with resistance, over a range of rod masses, clearly corroborating our working hypothesis.

**Figure 4:**
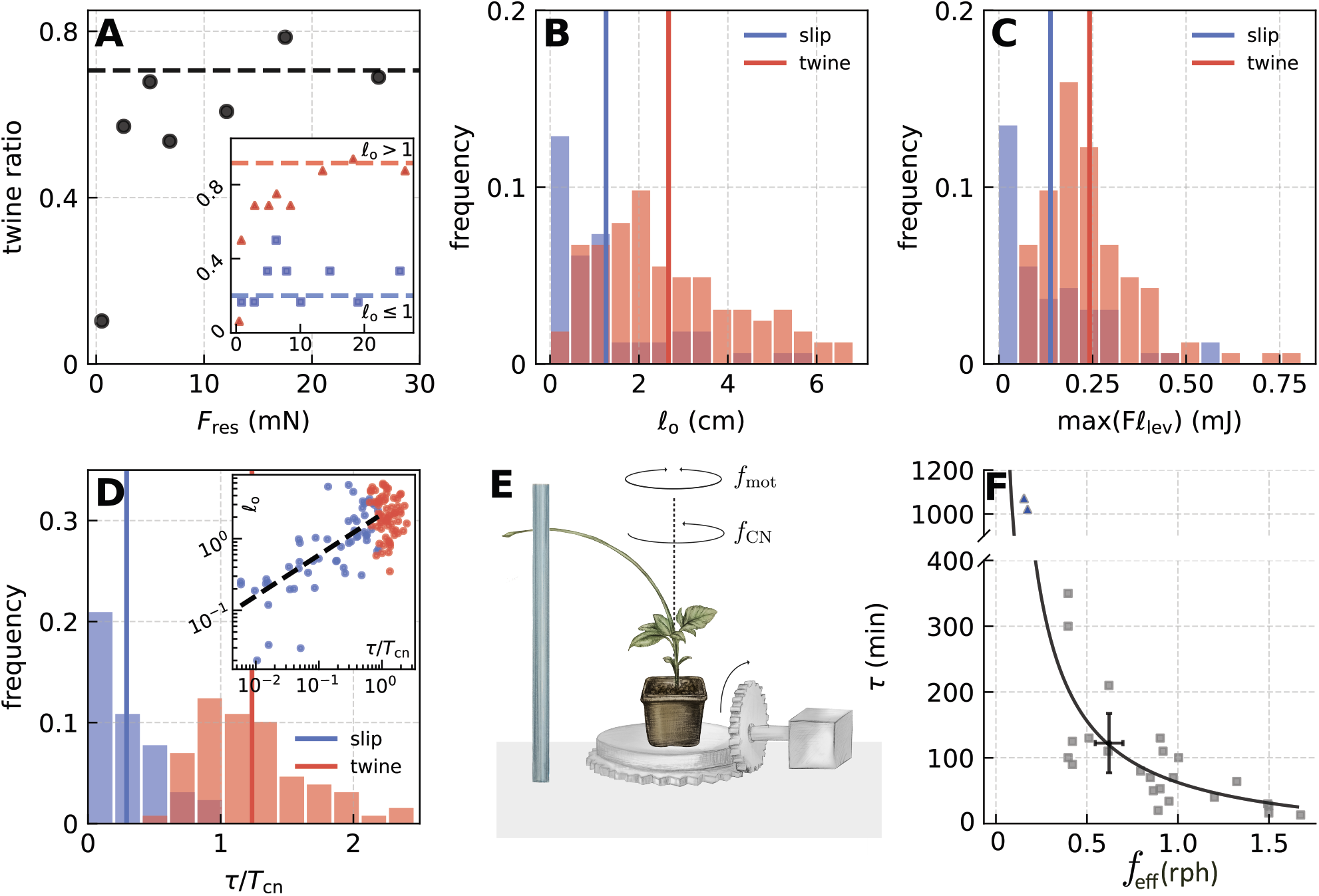
Twining initiation depends on support stability and grasp geometry. **A.** Dependence of the ratio of twining events (vs slip) on resistance force 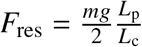 (*m* and *L*_p_ are the pendulum mass and length, and *L*_c_ is the distance of the contact point from the pendulum top (*20*)). Points show twining ratios for all events, binned by equal event counts (n=29 per point); the dashed line indicates the twining ratio for a fixed support (*F*_res_ → ∞, n = 17). (inset) Twining ratio for events with *ℓ*_o_ > 1 cm (red squares, n=16 per point) and *ℓ*_o_ ≤ 1 cm (blue squares, n=6 per point); dashed lines indicate the fixed-support limit. **B**. Distributions of *ℓ*_o_ prior to slip (blue, n=58) or twine (red, n=106) events. Vertical lines indicate mean values. Kruskal–Wallis p-value 7 × 10^−10^. **C**. Distributions of maximal torque max (*Fℓ*_lev_) for slip and twine events. Vertical lines indicate mean values. Kruskal–Wallis p-value 2 × 10^−6^. **D**. Distributions of contact time τ until slip or twine, normalized by the individual circumnutation period *T*_cn_. Vertical lines indicate mean values; Kruskal–Wallis p-value 8 × 10^−17^. (inset) Log–log plot of *ℓ*_o_ vs normalized contact time. Fitting slipping events yields *ℓ*_o_ = 0.87 (τ / *T*_cn_)^0.57^ (*R*^2^ = 0.53), consistent with Eq. 3. **E**. Schematic of the motorized setup. The plant is placed on a rotation stage in front of a static support (blue rod). The plant’s circumnutation is counter-clockwise (frequency *f*_cn_ = 1 / *T*_cn_), while the stage rotates clockwise or counter-clockwise with frequency *f*_mot_. **F**. Contact duration until twining initiation τ for motorized experiments with a fixed rod (grey squares), as a function of the effective rotation rate *f*_eff_ = *f*_cn_ ± *f*_mot_ (depending on motor direction). For comparison, the black cross denotes the mean contact duration measured in unperturbed experiments. Fitting the motorized data yields τ = 93/ *f*_eff_ − 31min (*R*^2^ = 0.53, 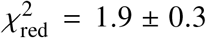). Blue triangles indicate events at very low frequency that did not twine within the observation time. Twining resumes once the motor is stopped.

We further reanalyzed the experiments by dividing them according to overshoot length (Fig. 1A), with *ℓ*_o_ =1 cm taken as an arbitrary threshold (inset Fig. 4A). For small overshoot, the twining ratio plateaued at approximately 0.2 and showed little dependence on resistance. For larger overshoot, twining probability rose sharply with increasing resistance. This separation demonstrates that the mechanical resistance alone is insufficient in the absence of adequate contact geometry, highlighting a geometrical requirement for grasp formation. To test this explicitly, we plot the distribution of overshoot length *ℓ*_o_ for both twining and slipping outcomes (Fig. 4B), finding a significant difference between the two with a Kruskal-Wallis pvalue of 7×10^−10^. Twining tends to occur for overshoots *ℓ*_o_ >1 cm, suggesting that, beyond support stability, plants initiate twining only when there is enough overshoot to make twining possible at all, similar to a reach-and-grasp task (*40, 41*), where the index finger needs to be positioned beyond an object for a hand to be able to grasp it.

To identify the signal that effectively triggers twining in climbing plants, we note that higher mechanical resistance, which is associated with an increased probability of twining, results in larger deformations of the plant shoot. It is widely accepted that plants perceive and respond to mechanical stimuli across multiple organizational scales (*28, 42–49*), from single cells to whole organs, and that these responses shape overall plant architecture. It can therefore be expected that, in climbing plants, increased shoot deformation may encode the mechanical signal that initiates twining.

To examine this hypothesis, since shoot deformations cannot be directly quantified in our setup, we take advantage of the moment equilibrium in Eq. 2 (Fig. 3E), which states that the internal moment due to bending *M*_in_ =*B*(*κ*−*κ*_0_)is equivalent to the external moment *M*_ex_ =*Fℓ*_lev_. The latter quantity can be measured directly. Fig. 4C shows the distribution of maximal external moments for each force trajectory, according to the different outcomes, revealing a significant difference between twining and slipping events, with a p-value 2×10^−6^ (Kruskal-Wallis 2-distribution test, see Methods). These distributions are consistent with a threshold deformation criterion, indicating that twining corresponds to a mechanical transition occurring when the stem reaches a critical load-dependent deformation set by support stability.

Having understood the structural and geometric component, next we consider the timescale of the problem, namely the time until slipping or twining occurs. Fig. 4D shows the distribution of duration of the contact between plant and rod normalized by the circumnutation time of the specific plant, τ/*T*_cn_, color-coded by twining and slipping outcomes. Experiments fall into two distinct domains, where contact durations below roughly one circumnutation period tend to lead to slipping, while longer contact durations, τ/*T*_cn_ ≳1, tend to end in twining.

Longer contact durations allow greater force accumulation and deformation. While it is clear that contact duration is governed by the circumnutation period *T*_cn_, it is also modulated according to the geometry of the contact point. Namely, longer overshoots *ℓ*_o_ mean that potential slipping takes longer, following, the relation

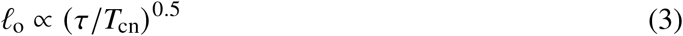

(see theoretical derivation in the SM). This is corroborated in the inset of Fig. 4D. Thus, larger overshoots delay detachment, increasing the likelihood that the system reaches the twining threshold. Longer overshoots are also stiffer at the contact point (Fig. S5B), further slowing down the slipping time.

Lastly, we tested the hypothesis that circumnutation period governs the time required to reach the threshold torque or deformation that triggers twining. We artificially changed the circumnutation period (described by the frequency *f*_cn_ =1/*T*_cn_) with a rotating stage with frequency *f*_mot_ which can rotate in the same or opposite direction of CNs (Fig. 4E). Fig. 4F shows the contact time until twining initiation, as a function of effective rotation frequency *f*_eff_ =*f*_cn_ ±*f*_mot_. Faster effective rotations led to faster twining initiation, even within minutes - significantly faster than natural twining times. Slower effective rotations, on the other hand, delayed twining initiation; in extreme cases, where rotation rate was less than 0.25 rph, twining did not occur for several hours, or did not occur at all. These cases are represented by blue triangles, where contact time is equal to the end of recording time (example in Movie S4). Twining resumed once the motor was stopped, restoring the natural force accumulation dynamics.

## Discussion

The results presented here show that climbing plants perform a form of active mechanical sensing, where the capacity to distinguish and select suitable supports emerges from the coupling between self-generated dynamics and intrinsic material properties. As in whisking in mammals, antennal probing in insects, or flagellar beating in microorganisms, self-generated motion is used to actively probe the environment. Unlike these systems, however, the sensing here emerges from the continuously growing organism itself rather than from a specialized sensory organ. By modeling the shoot as a slender active rod, the absence of centralized control can be understood in physical terms: the process of environmental discrimination is effectively implemented through the plant’s morphology. In this framework, the system represents a biological realization of morphological computation, where the tapered geometry and spatial variation of the Young’s modulus function as a distributed mechanical sensor, encoding the conditions for twining directly through the underlying physics.

By reducing the interaction between a circumnutating shoot and its environment to a minimal model of a rotating cantilever governed by torque balance, we identify the physical constraints that underlie this decentralized behavior. This framework reveals that the structural properties of the shoot are functionally organized for this sensing process. Near the apex, higher compliance and reduced diameter promote slipping, preventing attachment where the lever arm is too short to establish a grasp. Further from the tip, the increasing stiffness allows the system to reach a critical bending moment, which corresponds to a characteristic stem deformation. This state likely serves as the mechanical trigger for twining initiation (*47*), aligning with evidence that organ-level mechanical strain in plants activates signaling cascades associated with thigmotropic responses. In this way, the physical architecture of the stem simultaneously filters for support stability and geometric overshoot, such that a climbing plant may undergo several successive contact events before twining is successfully initiated, with each encounter constituting a new mechanical probe. Attachment occurs only when the stem–support configuration satisfies the embodied mechanical criteria encoded by the stem’s morphology and elasticity.

Experiments with altered rotation periods decouple the role of time-dependent growth from the sensing dynamics. The observation that twining initiation can be accelerated or suppressed by modulating the circumnutation frequency indicates that these oscillations define a probing timescale. This distinguishes stem twiners from local touch-responsive systems such as tendrils: here, sensing occurs through organ-level mechanical integration, with the plant responding to the global torque state rather than to a discrete contact.

Broadly, these findings expand the functional scope of circumnutations, showing that they are not limited to a passive search for supports through sweeping dynamics, but represent a fundamental mechanism for active mechanical sensing. Rather than merely discovering the presence of an object, these oscillations enable the plant to actively evaluate the physical suitability of its environment. This perspective complements recent studies in non-climbing species which have begun to reframe circumnutations as functionally important rather than incidental growth dynamics (*50, 51*). By demonstrating how a decentralized system achieves robust environmental selection through embodied dynamics, this work provides a blueprint for the design of intelligent matter and autonomous soft robotics. Our results suggest that future engineered systems could achieve complex navigation and interaction by tuning the morpho-elastic properties of their appendages, allowing the material body to sense and act without the need for sensors or centralized computation.

## Materials and Methods

### Growth conditions

Climbing bean plants (*Phaseolus vulgaris*, Helda strain) were grown in lab conditions, seeds were germinated on moist cotton in darkness at 24^°^C, then transferred soil pots and grown under fixed illumination with a 16 h light cycle. Plants were fertilized once with 2 ml of 2% 20-20-20 N–P–K fertilizer. Experiments were performed on 2-4-week-old plants whose height, measured as the highest point on the bent stem, was between 20 and 40 cm. The horizontal searcher stem extending beyond the pot was typically 10-20 cm long. Experiments were conducted under constant LED illumination of 8 ± 2 W m^−2^ (*20*). We also present the growth rate along the stem in Fig. S11.

### Experimental setup

Forces exerted by the plant stem were measured using a pendulum-based setup. A lightweight rod was suspended as a pendulum near the plant such that the stem contacted the rod during circumnutation. Rod deflection upon contact was recorded using two orthogonally positioned cameras (top and side views). The position of the stem and the resulting rod displacement were tracked over time and used to infer force trajectories. Measurements were performed using pendulum rods with masses between 0.5 and 3 g. As described in (*20*), different pendulum masses were used to extend the measurable force range of the setup while maintaining force sensitivity over different loading conditions. Plants were positioned so that the stem encountered the rod naturally during circumnutation. Because a given plant may undergo several successive contact events before twining is successfully initiated, each contact event was treated as an independent observation (*20*) and classified according to whether it resulted in twining or slipping (see below).

### Image analysis

Tracking of the tip of the pendulum rod and the position of contact between the stem and the rod described in (*20*) (including script). Tracked movements used for force measurements. We extracted the stem centerline (Fig. S12) using thresholding and masking functions (*52*), skeletonize (*53*), smoothing,interpolation and nearest-neighbor ordering (*54*). Script available on (*55*). Converting measured pixel lengths to physical lengths was done using a reference object of known length (different per view) (*20*).

### Force trajectory extraction and normalization

Plants were placed within a custom-built force measurement setup developed in our laboratory (*20*). The bean stem interacts with a physical pendulum. Based on the measured deflection angle of the pendulum, together with known system parameters, we calculated the horizontal force exerted by the plant using a torque equilibrium equation for up to one CN period, as detailed in (*20*) (example in Movie S1 of (*20*)). We filtered events where the stem only contacts the rod from below, as there is nearly no normal force and so not relevant for our analysis. Each force trajectory is smoothed using a moving window average, and first half of the CN period is fitted to a sine function 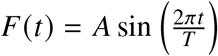 with fitting parameters A and T. We normalize trajectories, dividing force values by A, and elapsed time by T.

### Physical model parameter measurements

Here we detail the measurement of parameters in Table 1. **Radius:** At the end of the experiment, the diameter of the stem was measured at different distances from the stem tip for *L* − *s* ∈ {5, 10, 15, 20, 25, 35, 45} cm. Specifically, for each plant, 5 − cm stem segments were sequentially excised starting from the apical tip, and continuing up to *s* = *L* − 50 cm. Stem diameter was measured using a digital caliper or with image analysis after the experiment, yielding the stem radius. **Young’s modulus:** For each 5 − cm segment, we measured the Young’s modulus *E* by subjecting the segment to a three-point bending test, in which incremental weights were applied near the midpoint of the segment with a suspended string to induce bending (Fig. S13). The stem was placed between two supports aligned with a Nikon D7500 camera, with a Tamaron 24-70 mm f/2.8 Di g2 macro lens attached. 3D printed setup .stl files are available at (*55*). Further details in the SM. **Circumnutation times:** Before force measurements were performed, each plant was tracked while performing free CN rotations for one or more periods, extracting each plant’s mean CN period. **Maximal curvature** *κ*_**max**_ : Analyzing side view images of the stem during the natural CN period (before contact with the support). We extracted the centerline using image analysis, and calculated the local angle and curvature (Fig. S14).**Overshoot and lever lengths** *ℓ*_**o**_ **and** *ℓ*_**lev**_ : The overshoot length was measured from the top view image (or side view, if top view was irrelevant) at the time of contact. To extract the lever length, we first need to assess the length of the whole stem

*L*. We do not have a direct measurement of this, we therefore add the length of the stem caught in the side view image *l*_skel_, and assess the missing length from the image cutoff to the plant base by subtracting the y-component of the image *y*_skel_ from the total vertical height of the bean plant *h* along the y-axis, measured at the beginning of the experiment:*l*_base_ ≈ *h* −*y*_skel_ (Fig. S12E). Thus *L*=*l*_skel_ +*l*_base_. Finally, the lever length is equal to *ℓ*_lev_ =*L*−*ℓ*_o_.

### Determining twining onset based on stem angle

Twining initiation is a gradual rather than discrete process. To quantitatively identify its onset, we therefore defined a threshold based on the angle between the stem and the support. Based on visual observations of the onset of stem wrapping, we chose a threshold angle of approximately 45^°^ with respect to the pendulum (see Movie S1 for an example of a twining event). The estimated initiation times were largely insensitive to the precise threshold choice; using a threshold of 50^°^ instead of 45^°^ produced nearly identical results for the vast majority of events. To determine the moment twining began, we segmented images of the stem from the side view during contact with the support using the Interekt software (*56*), and extracted the angle near the contact position with the support. In cases where Interekt failed, we tracked two points on the stem on either side of the contact point from the side view (tracking script in (*20*)).

### Statistical analysis

We distinguished statistical difference between distributions using Kruskal–Wallis test for 2 groups (effectively a Mann-Whitney U test) as post hoc testing. Statistical tests were done using the ‘Scipy’ Python package (*54*).

### Motorized stage

We built a rotating stage using a standard 0.5 rpm, 12 V spur gear DC motor (SparkFun Electronics) powered by a 12V supply. Because this rotation rate exceeds the natural circumnutation of the plants, the motor speed was reduced through a gear system. A 1 : 60 ratio worm gear-box (*57*) was coupled to the motor via an aluminum alloy flange (secured with M3 screws), followed by an additional gear set with a ratio of 1 : 3, yielding a final rotation rate of 1 : 180 of the motor’s original speed. A plate mounted atop the last gear provided support for the plant pot (*55*). Experiments were recorded via timelapse for up to 10 hours from top and side view angles, capturing slip and twine events, with some plants exhibiting multiple events. Motor control was implemented using Arduino UNO/MEGA micro-controllers, with an H-bridge modulating the power supply in response to pulse width modulated (PWM) signals from the Arduino. Because the motor rotation rate exhibited a nonlinear dependence on the input PWM signal, the final rotation speeds were determined empirically by tracking a reference point on the stage using the top-view camera during the experiment. Prior to initiating contact, the plant’s natural CN period *T*_cn_ was recorded over at least one full rotation. The plant was then positioned near a stable support, and the motor activated at a predetermined rate set via the Arduino-controlled PWM. At low effective rotation rates (i.e., motor rotation rate nearly equaled the plant CN rate), a brief period of contact with naturally progressing circumnutations was allowed (pre-straining) to accommodate fluctuations in CN and prevent premature detachment from the support.

### 3D printing

All 3D printed objects were done with a Sindoh 3D WOX DP200 printer with ELEGOO, 1.75 mm diameter PLA filament. Object files can be in the Zenodo repository (*55*).

### Numerical simulations

The simulations were performed using the Elastica numerical solver (*24, 36*) presented in (*39*). Simulation software available in (*58*). The solver assumes a separation of timescales between slow growth-driven plant responses and fast mechanical relaxation, allowing us to decouple the two processes in a quasi-static manner (*38*). Simulation parameters were chosen to remain of the same order of magnitude as the experimental measurements (Table 1), rather than to reproduce individual plants quantitatively. In particular, shorter organ lengths substantially reduce computation time, since the relaxation time step is set by the elastic shear-wave travel time across a discretization length and relaxation requires many wave-travel times across the organ (*39*). Young’s modulus values were similarly selected empirically to provide computationally efficient simulations.

#### Shape

At each arc length *s* along the centerline of the simulated stem we define the position vector ***r***(*s, t*), which provides the 3D description of the rod at time *t*. We also define a local orthonormal material frame {**d**_1_ (*s, t*),**d**_2_ (*s, t*),**d**_3_ (*s, t*)}, where **d**_1_ and **d**_2_ span the cross-section of the organ, and in a shearless and inextensible system **d**_3_ coincides with the tangent of the centerline. The local curvature vector ***κ***(*s, t*) (or Darboux vector) of the center-line is defined through the the relation 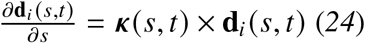. The components of curvature projected along the principal vectors of the material frame (***κ*** = *κ*_1_ **d**_1_ +*κ*_2_ **d**_2_ +*κ*_3_ **d**_3_) coincide with bending (*κ*_1_, *κ*_2_) and twist (*κ*_3_) strains in the material frame. Introducing elastic stretch may lead to a difference between the arc-length configuration *s* and the stress-free configuration *s*, described by the local stretch (*24*) 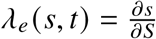. Shear may lead to an incongruity between **d**_3_ and the tangent of the center-line, and the local deviation is described by the translation vector 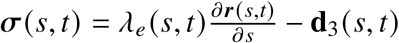.

#### Model for circumnutations

Here, growth is taken to be implicit, such that the organ does not increase its length but remodels it intrinsic curvature 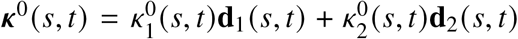.

Note that 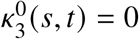 such that no intrinsic twist is assumed. To emulate a circumnutating stem we follow (*59*) and model the dynamics of the intrinsic curvature vector of the stem as (omitting the *s, t*dependencies for brevity):

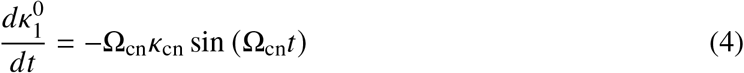

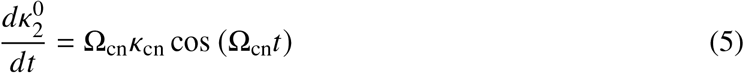

where Ω_cn_ = 2*π*/*T*_cn_ = 2*π f*_cn_ and *κ*_cn_ are the circumnutation angular velocity and curvature correspondingly. By choosing the initial intrinsic curvature to be ***κ***_0_(*s*, 0)=*κ*_cn_**d**_1_(*s*, 0), Eqs. 4,5 give:

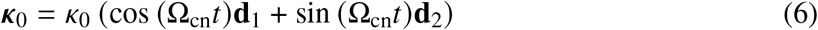

such that the intrinsic shape of the organ is always a constant curvature *κ*_0_, which rotates in a circular motion without twisting the material frame (see Movie S2).

#### Mechanics

Mechanical equilibrium is achieved at each time step when the time-independent Cosserat rod equations are fulfilled along the center-line (*24*) (omitting the *s, t* dependencies for brevity):

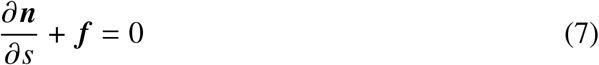

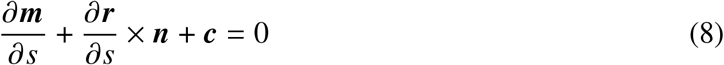

where ***n*** are the internal contact forces, ***τ*** are the internal torques or bending moments, ***f*** are the external forces per unit length and ***c*** are the external couples per unit length, all functions of *s* and *t*. The full elastic relaxation dynamics between growth time steps are based on the Cosserat model with a dissipation mechanism, as described in (*24*). We follow (*24*) and choose a constitutive model that assumes linear elasticity, namely the internal contact forces are linearly related to the shear and stretch of the centerline, and the internal torques are linearly related to the bending and twisting:

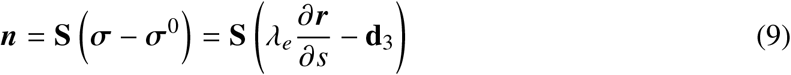

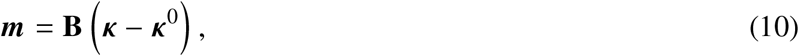

where **S** and **B** are stiffness matrices which are related to the cross-sectional area and the second moment of inertia such that **S** = *E πR*^2^ · diag(8/9, 8/9, 1) and **B** = *E πR*^4^ · diag(1/4, 1/4, 1/6), expressed in the local material frame (*24*). In our simulations we use *E* ∈ {10, 50, 100} MPa. We also assume volume conservation, varying the local radius of the organ in order to compensate for the elastic stretch 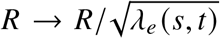. The contact between the obstacle and the organ is modeled using a spring-like force with a spring constant of *k*_spring_ = 100kg/s^2^, and we assume both friction and friction-less contact. The dissipation mechanism of the elastic relaxation is modeled using a friction coefficient of *ν* = 0.01 kg/sec. For more details see (*24, 39*).

## Acknowledgements

We thank Roni Kempinski, Alessia Perilli, Mathieu Riviere and Agueda de la Vega, for consulting, assisting and troubleshooting. We thank Karen Marron, the best scientific editor in the universe, and Barak Hadad, who designed beautiful setup illustrations inspired by Charles Darwin.

## Funding

Y.M. acknowledges support from the Israel Science Foundation Research Grant (ISF) no. 2307/22, ERC grant GROWsmart 101165101, and European Union’s Horizon 2020 research and innovation program under Grant Agreement No. 824074 (GrowBot). A.O. acknowledges support from the Colton Foundation scholarship. A.P. acknowledges funding from the Gatsby Charitable Foundation grant GAT3395/PR4B, and the Herchel Smith Postdoctoral Fellowship Award.

## Competing interests

None declared.

## Author contributions

A.O: methodology, experimental design, carried out all experiments, experimental data curation; A.P: developed simulation framework, carried out simulations, simulation data curation; A.O, A.P and Y.M: formal analysis, conceptualization and investigation, visualization, writing original draft, review, and editing, Y.M. funding acquisition, supervision, and resources.

## Data, code and materials availability

All data and code needed to evaluate and reproduce the results in the paper are present in the paper and/or the Supplementary Materials. This study did not generate new materials.

Datasets and analysis scripts used to generate trajectories are openly available via Zenodo: https://zenodo.org/uploads/18865114 (*55*). Includes an Excel spreadsheet containing event data including dates, radius, rod mass and length, contact initiation, and experimental data for Young’s modulus extraction. A README file details folder contents.

The Github repository https://github.com/merozlab/ohad-2026-cn-mech-twine.git includes Python scripts for importing data and calculating force trajectories, curvature, Young’s modulus, and fitting and generating figures.

## Supplementary materials

References *(7-60)*

Supplementary Text

Figs. S1 – S14

Movies S1 – S4

## Notes

https://doi.org/10.5281/zenodo.18865113

### Competing Interest Statement

The authors have declared no competing interest.

### Summary of Updates

Changes after reviewer comments: updated abstract; updated introduction; added details for simulation parameter assumptions and value choices; added video exhibiting circumnutation and twining motions; updated discussion; corrections to Table 1 caption; corrections in captions of Figures 1-4; minor corrections to Figures 1-4, added detail to Materials and Methods; Github link added in data and code availability; added diagram Figure S1, schematics Figures S8 and S9, growth rate S11, movies S1, S2 (previous movies are S3,S4)

