## Supplementary for "Embodied mechanical sensing through self-generated motion in climbing plants"

[†]These authors contributed equally to this work.

email address [*]Corresponding author. Email:

**This PDF file includes:**

Supplementary Text

Figures S1 – S14

Captions for Movies S1 – S4

**Other Supplementary Materials for this manuscript:**

Movies S1 – S4

### Supplementary Text

#### Estimation of Young's Modulus

Deflection profiles obtained through image analysis from the weights applied to each stem segment were analyzed using Euler–Bernoulli beam theory with the corresponding boundary conditions, namely an asymmetrically loaded beam with two free end supports (*60*):

$$\begin{aligned} w &= \frac{PbL_{\mathrm{b}}x(L_{\mathrm{b}}^2 - (bL_{\mathrm{b}})^2 - x^2)}{6L_{\mathrm{b}}EI} + \frac{P(x - aL_{\mathrm{b}})^3}{6EI}, \ x > aL_{\mathrm{b}} \\ w &= \frac{PbL_{\mathrm{b}}x(L_{\mathrm{b}}^2 - (bL_{\mathrm{b}})^2 - x^2)}{6L_{\mathrm{b}}EI}, \ x <= aL_{\mathrm{b}} \end{aligned} \tag{S1}$$

Each loaded segment was compared to the free segment shape. Thereafter the Young's modulus (effective value per section, as stem is not homogeneous) was extracted by fitting analytical bending solutions (Eq.S S1) to the curvature data and averaged over several loads to reduce measurement uncertainty (Fig. S13).

where $P$ - external load in grams, $a$ - distance of loading point from first base, $b$ - distance of loading point from second base, $L_{\mathrm{b}}$ - distance between base positions, $x$ - position from segment end, $E$ - Young's modulus, $I$ - second moment of inertia, in this case assumed circular cross section.

We note that, in many experiments, some measurements of radius or Young's modulus were unavailable due to experimental limitations, including measurement error and stem damage or dehydration following removal from the support (typically after several hours of twining). Consequently, the data were non-uniform across plants, motivating the use of mean and fitted values for the physical parameters.

#### Slip timing analysis

As stem stiffness near the tip decreases rapidly near the tip, we estimate the time it takes the stem to slip off of support due to large curvature of the organ using experimental relations of the radius, Young's modulus and force.

**Stem setup** We consider an Euler–Bernoulli cantilever beam of length $L$, clamped at $x = 0$ and free at $x = L$. A transverse force is applied at a point located a distance $a$ from the tip, i.e. at

$$x_c = L - a.$$

We define the coordinate

$$s = L - x,$$

which measures distance from the tip toward the base. Thus the tip corresponds to $s = 0$ and the base to $s = L$.

**Stem tapering and stiffness** Experiments indicate that the Young's modulus varies near the tip as

$$E(s) = E_{\text{tip}} + \alpha_E s^2.$$

The stem radius increases approximately linearly with distance from the tip, so the second moment of area varies approximately as

$$I(s) = I_{\text{tip}} + \alpha_I s^4.$$

The bending stiffness is therefore

$$B(s) = E(s)\, I(s).$$

**Curvature** For a cantilever with a point force $F$ applied at $s = a$, the bending moment for $s \geq a$ is

$$M(s) = F(s - a).$$

Euler–Bernoulli beam theory gives the curvature:

$$\kappa(s) = \frac{M(s)}{E(s)I(s)}, \qquad \frac{d\theta}{ds} = \kappa(s),$$

with $\theta$ the rotation angle.

**Tip angle** The tip angle equals the rotation at the point of load application:

$$\theta_{\text{tip}} = \theta(s = a) = \int_a^L \frac{F(s - a)}{E(s)I(s)}\, ds.$$

Substituting the expressions for $E(s)$ and $I(s)$ yields the exact integrand

$$\frac{F(s - a)}{\left(E_{\text{tip}} + \alpha_E s^2\right)\left(I_{\text{tip}} + \alpha_I s^4\right)}.$$

**Integrated Tip Angle** Integration yields the tip angle

$$\theta_{\text{tip}} = \int_a^L \frac{F(s - a)}{\left(E_{\text{tip}} + \alpha_E s^2\right)\left(I_{\text{tip}} + \alpha_I s^4\right)}\, ds.$$

**Approximations** Assuming low constant values near the tip we get an approximation:

$$E(s) \approx \alpha_E s^2, \qquad I(s) \approx \alpha_I s^4,$$

so the bending stiffness becomes

$$B(s) \approx \alpha_E \alpha_I s^6.$$

Thus,

$$\theta_{\text{tip}} \approx \frac{F}{\alpha_E \alpha_I} \int_a^L \left( \frac{1}{s^5} - \frac{a}{s^6} \right) ds = \frac{F}{\alpha_E \alpha_I} \left( \frac{-4}{s^4} - \frac{-5a}{s^5} \right)_a^L$$

$$= \frac{F}{\alpha_E \alpha_I} \left( \frac{-4}{L^4} - \frac{-4}{a^4} - \frac{-5a}{L^5} + \frac{-5a}{a^5} \right)$$

Approximating $a \ll L$ we get the relations:

$$\frac{1}{L^4} << \frac{1}{a^4}; \frac{a}{L^5} = \frac{a}{L} \frac{1}{L^4} << \frac{a}{L} \frac{1}{a^4} << \frac{1}{a^4}$$

and we simplify the result of the integral to:

$$\theta_{\text{tip}} \approx \frac{F}{20\, \alpha_E \alpha_I} \frac{1}{a^4}.$$

Experiments and simulations show that the force grows approximately sinusoidally. For short contact durations we approximate a linear rise:

$$F(t) \approx F_0 t.$$

Additionally, we find the force magnitude scales approximately linearly with the Young's modulus and parabolically with the contact distance from the tip. We model:

$$F \propto E, F \propto s^2$$

so

$$F(t, s) \approx F_0 t E(s) \approx F_0 t s^2$$

Substituting into the expression for $\theta_{\text{tip}}$:

$$\theta_{\text{tip}}(t, a) = \frac{F_0}{20\, \alpha_E \alpha_I} t\, a^2.$$

**Force equilibrium** Let $\theta_{\text{base}}$ be the base rotation. The bending at the point of contact relative to the base is

$$\Delta\theta = \theta_{\text{tip}} - \theta_{\text{base}}.$$

Force equilibrium perpendicular to the stem gives the normal force:

$$F_{\perp} = F \cos \Delta\theta.$$

The friction force is approximately:

$$F_{\text{fric}} = \mu N = \mu F \cos \Delta\theta.$$

The normal force pushes into the support while the tangential component pulls the stem:

$$F_{\|} = F \sin \Delta\theta.$$

Slipping occurs when

$$F_{\|} = F_{\perp},$$

i.e.

$$\tan \Delta\theta = \mu.$$

For friction coefficient $\mu$ of order 1 this yields a critical angle:

$$\Delta\theta_{\text{crit}} \approx \arctan(\mu) \approx \pi/4.$$

**Angle-dependent slip** Set the critical angle $\Delta\theta_{\text{crit}} = \pi/4$ and solve

$$\Delta\theta(t, a) = \frac{\pi}{4}.$$

Using the expression above,

$$\frac{F_0}{20\alpha_E \alpha_I}\, t_{45}(a)\, a^2 = \frac{\pi}{4}.$$

Thus the arrival time to reach 45° is

$$t_{45}(a) = \left(\frac{5\pi\, \alpha_E \alpha_I}{F_0}\right) a^2.$$

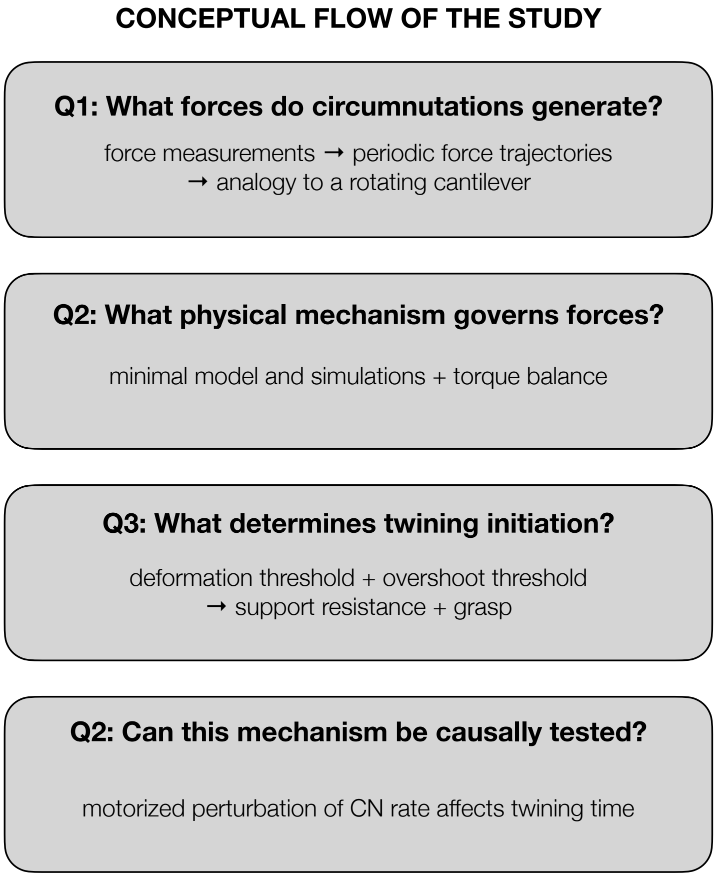


**Figure S1**: **Diagram summarizing the conceptual flow of the study and the logical progression of the questions addressed in this work.**

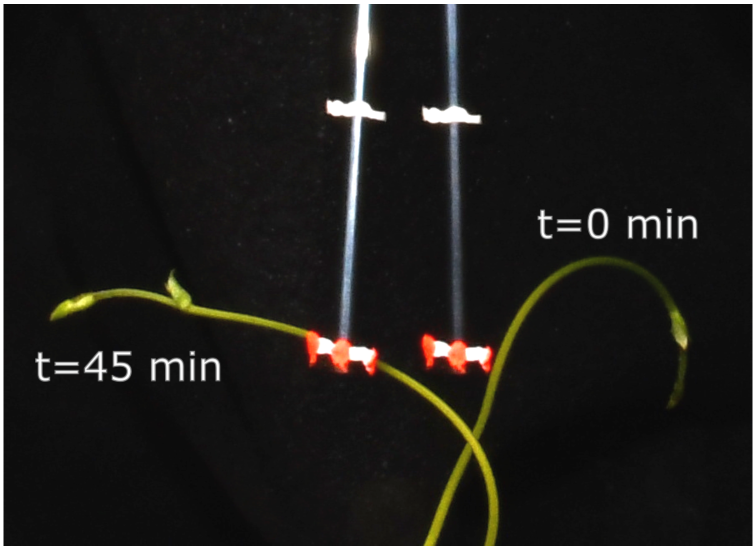


**Figure S2**: **Pendulum deflection snapshots.** Two snapshots of a circumnutating stem interacting with the pendulum rod (side view). First snapshot right before contact at t=0 (no deflection), and second snapshot after t=45 min of contact (rod deflected left).

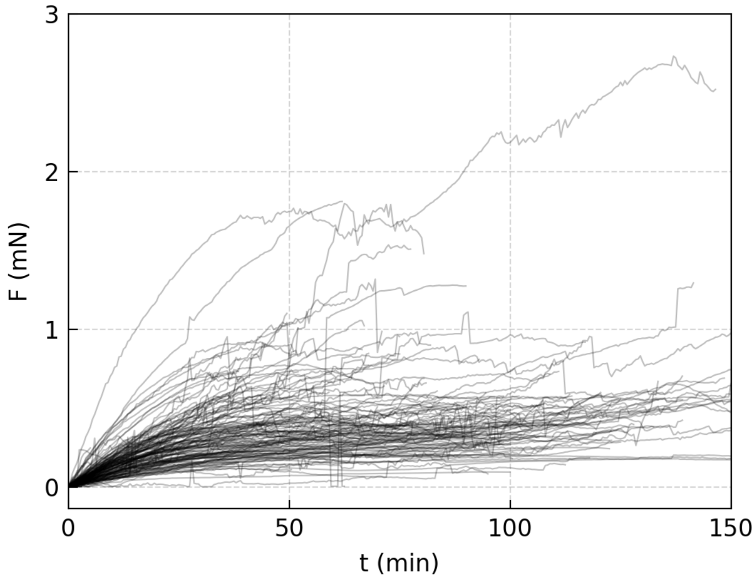


**Figure S3**: **Force trajectories.** Measured force trajectories extracted using pendulum force setup(n=164).

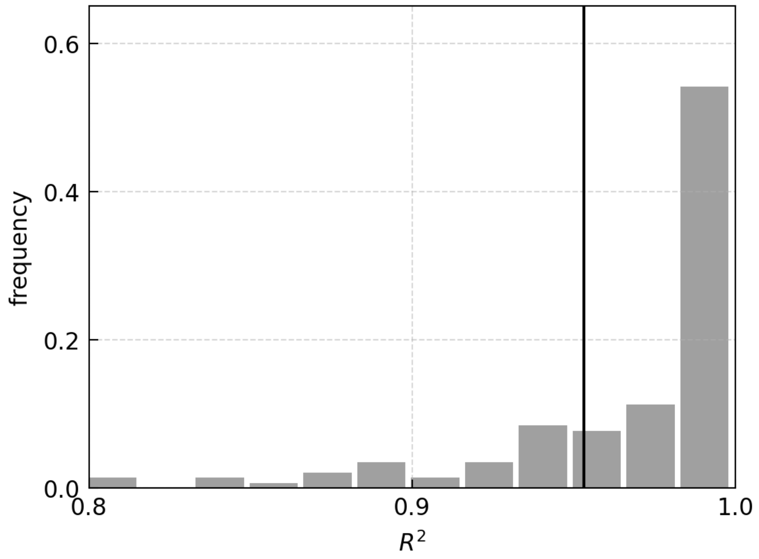


**Figure S4**: **Fit value of sine to measured force trajectories.** Distribution of $R^2$ values for fit to force trajectories shown in Fig. 1C.

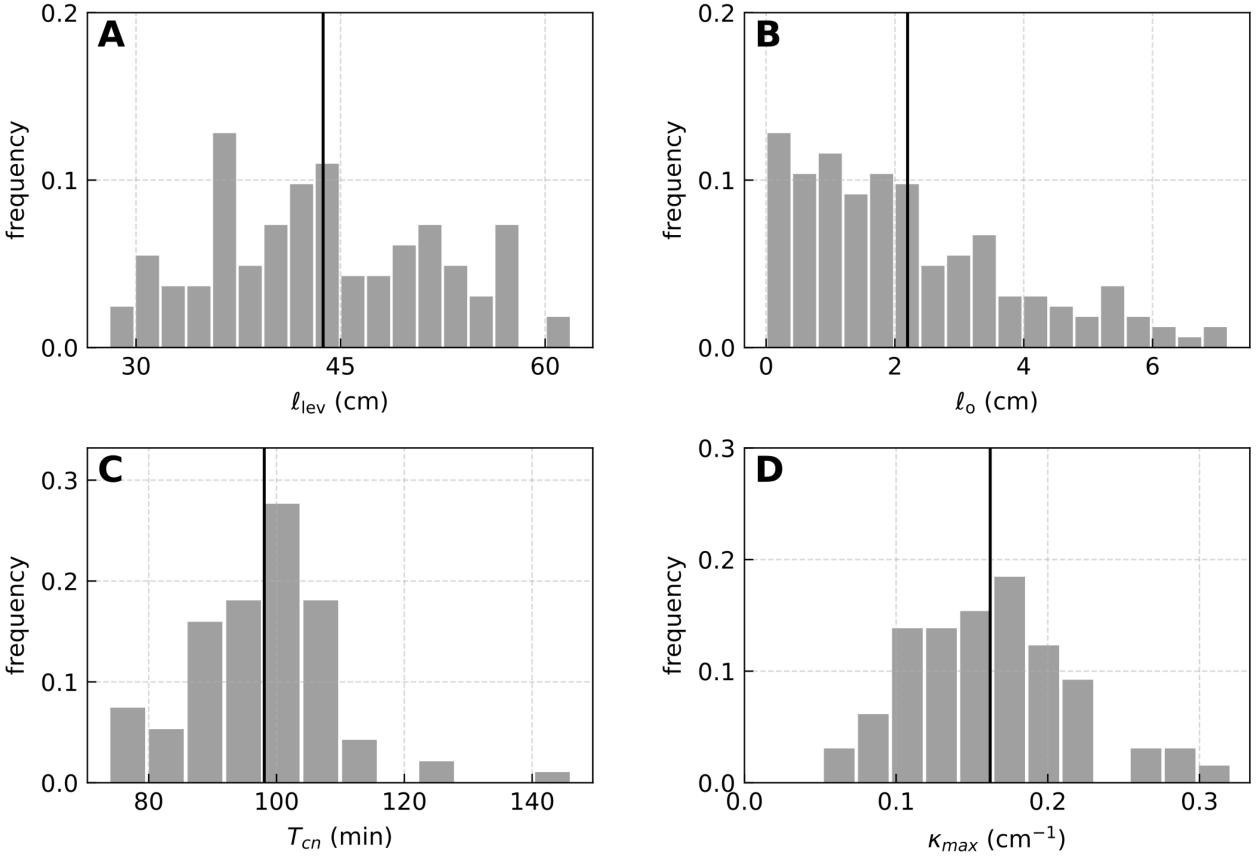


**Figure S5**: **Distributions of measured parameters. A.** Distribution of $\ell_{\text{lev}}$ values for pendulum events estimated from side view images(see methods). The mean lever value is: 42 ± 6 cm. **B.** Distribution of $\ell_{\text{o}}$ values (n=106) for pendulum events estimated from side view images(see methods). The mean overshoot value is $\ell_{\text{o}} = 2.1 \pm 1.6$ cm **C.** Distribution of mean circumnutation period $T_{\text{cn}}$ per plant (n=94), measured before contact. Mean value is 98 ± 11 min. **D.** Distribution of maximal curvature of the stem $\kappa_{\text{max}}$ before contact (n=65), extracted by fitting side images of the stem to a circle (Fig. S14 in SM). Mean value is 0.162 ± 0.054 $\text{cm}^{-1}$.

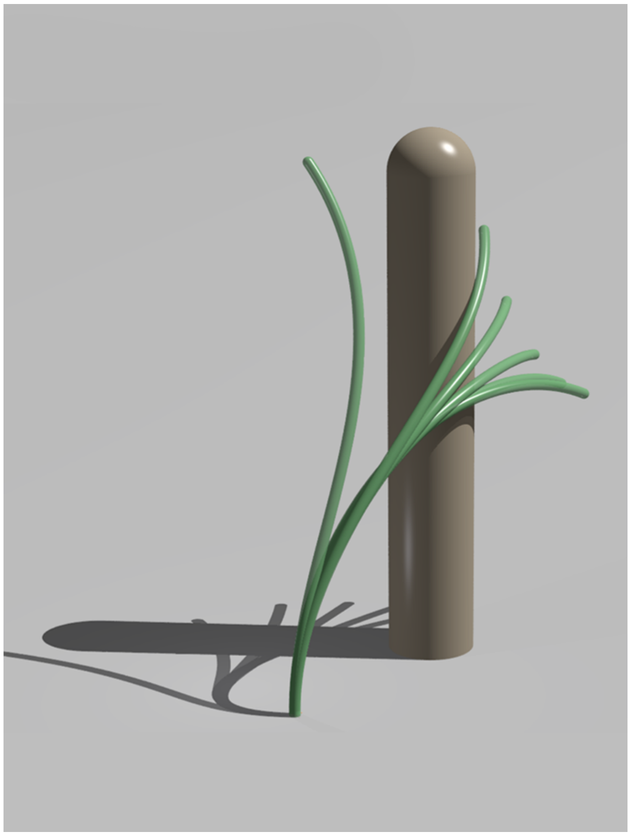

**Figure S6**: **Graphic rendering of a typical simulation.** The stem is modeled as a morpho-elastic rod of length $L = 5$ cm, with a constant radius $R$, Young's modulus $E$, and intrinsic curvature $\kappa_0$ throughout. It is clamped at its base and free at its apex. Without changing its length, the rod remodels its intrinsic curvature quasi-statically according to Eq. 6, following circular circumnutations with a period $T_{\text{cn}}$, anti-clockwise. Values are experimentally informed, shown in Table 1. The rod then comes in contact with a cylindrical obstacle of radius $R_{\text{obs}} = 0.4$ cm. Snapshots of the simulation are shown at times $\frac{t}{T_{\text{cn}}} \in \{0, \frac{1}{12}, \frac{2}{12}, \frac{3}{12}, \frac{4}{12}, \frac{5}{12}\}$ (full video in Movie S3). The point of contact moves closer to the apex with time, resulting in a slip.

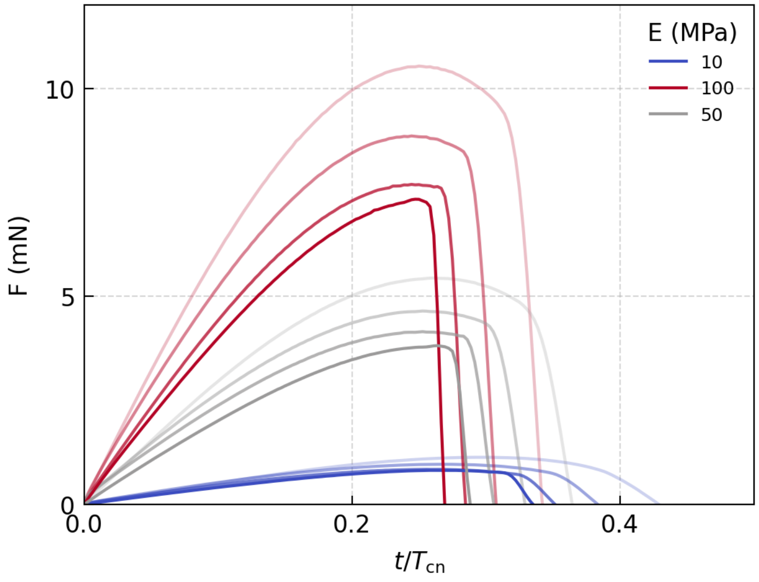


**Figure S7**: **All simulation force trajectories from Fig 3A, C, E.** Simulated forces with different Young's modulus (color label) and contact distances (lower intensity corresponds to larger $\ell_\mathrm{o}$ or smaller $\ell_\mathrm{lev}$). Shown together for further comparison.

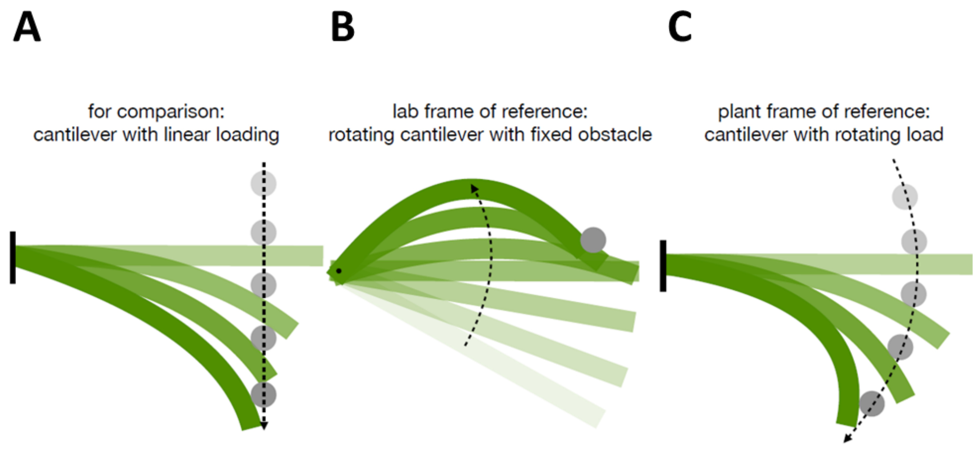


**Figure S8**: **Schematic of cantilever settings**. **A.** Classic cantilever loaded by obstacle moving linearly with speed $v$, where force follows $F = kvt$ over time, with $k$ the elastic spring constant of the cantilever. **B.** A simplified picture of the circumnutating plant encountering a support (projected on the x-y plane, top view) can be thought of as a rotating cantilever encountering a fixed load. In lab frame of reference. **C.** Shifting to the frame of reference of the plant (cantilever) this system is just a clamped cantilever with rotating load, thus yielding $F = kv \sin 2\pi t/T$, where $T$ is the period of rotation.

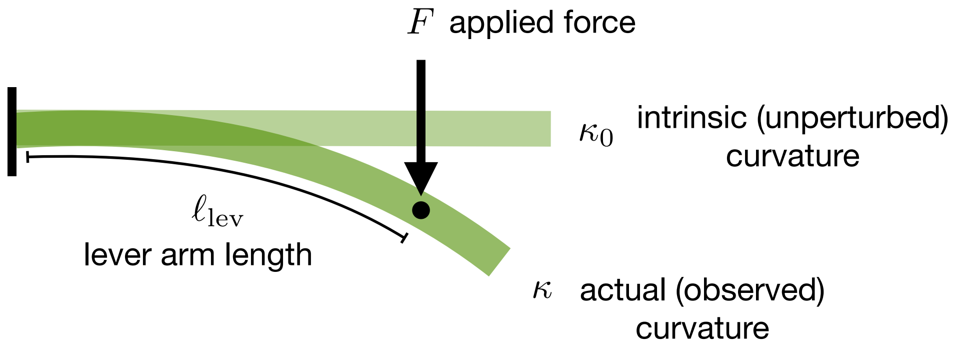


**Figure S9**: **Schematic illustration of the torque balance underlying Eq. 2.** A force F applied at a distance $\ell_{\text{lev}}$ from the clamped base generates an external moment $M_{\text{ext}} = F\ell_{\text{lev}}$. This is balanced by the internal bending moment $M_{\text{int}} = EI(\kappa - \kappa_0)$, arising from the deviation of the observed stem curvature $\kappa$ from its intrinsic (unperturbed) curvature $\kappa_0$. Equating the external and internal moments yields Eq. 2, relating the applied force to the stem's material properties, geometry, and deformation.

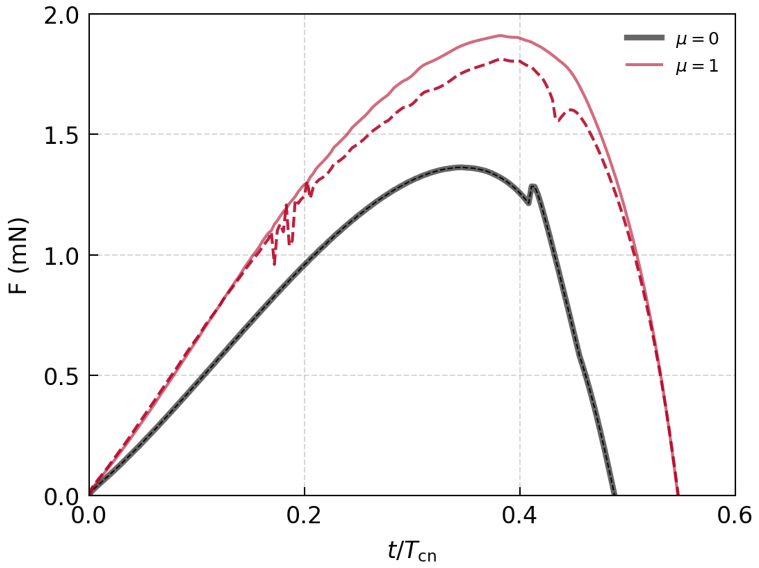


**Figure S10**: **Effect of friction in force simulations, comparing full force and 2D projection of pendulum measurement.** Force trajectories of the force (smooth line) with the support with friction ($\mu = 1$, red) and without friction ($\mu = 0$, black), for simulation trajectories with Young's modulus $E = 10$ MPa. Dashed line represents 2D projection of the force, equivalent to what is measured in the experimental setup. The difference between the full force and the planar force increases in the presence of friction.

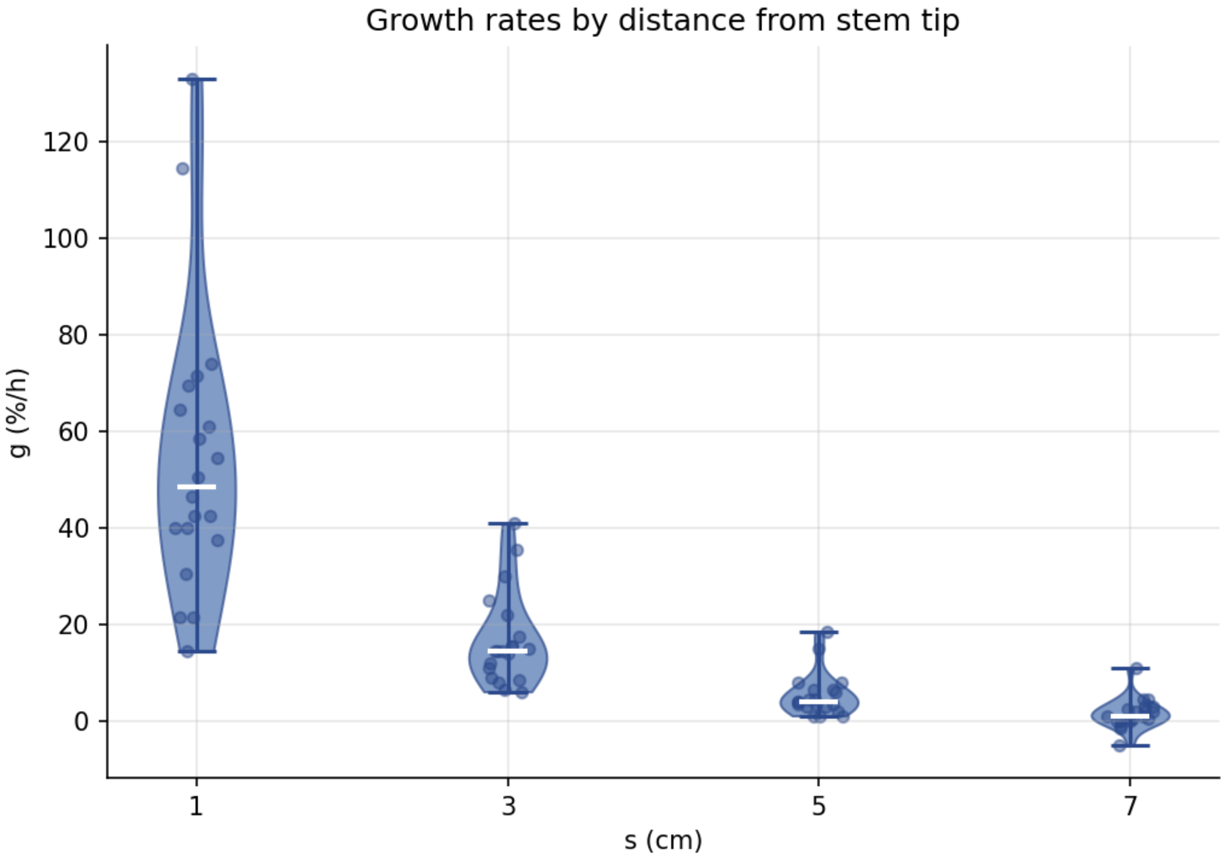


**Figure S11**: **Measured relative elongation along the stem.** Paint marks were applied at equal spacing along the stem at different distances from the tip. After several hours, segment elongation was measured relative to the initial segment length. Elongation decreased with distance from the tip and approached zero at approximately 8 cm, marking the end of the growth zone.

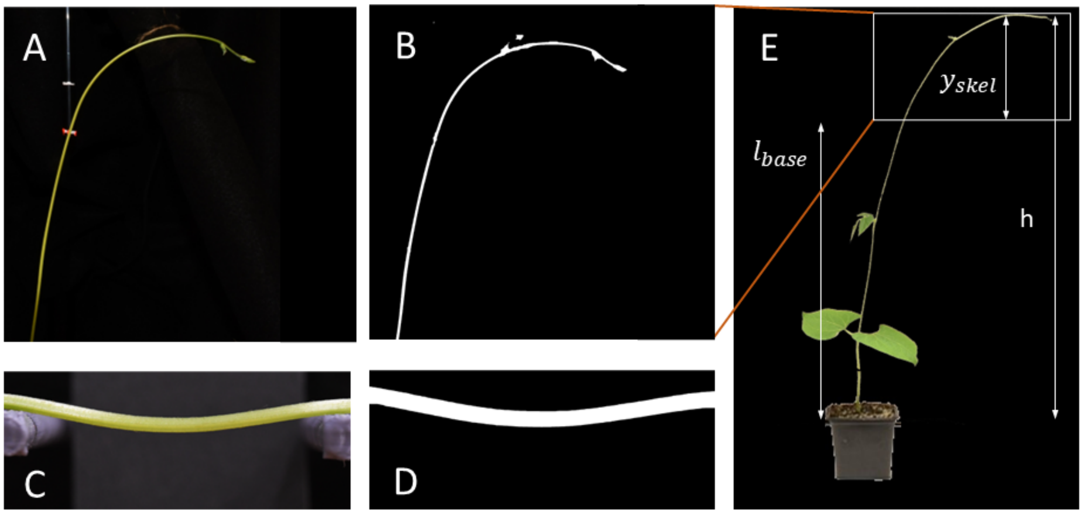


**Figure S12**: **Stem imaging and lengths A.** Stem side view image from force extraction setup. **B.** Image in A. after filter and smoothing. **C.** Stem side view image for 3-point bending measurement after force measurements were complete. **D.** Image in C after filter and smoothing. For all filters we applied a hsv filter, used threshold masking and smoothing functions to segment the stem from the background. These were later used to extract the centerline of the stem for further analysis(S14). **E.** Filtered side view image of full plant stem, showing the measured parameters $h$, $y_{skel}$ and $l_{\text{base}}$, using image B. as a reference for how the lengths were determined.

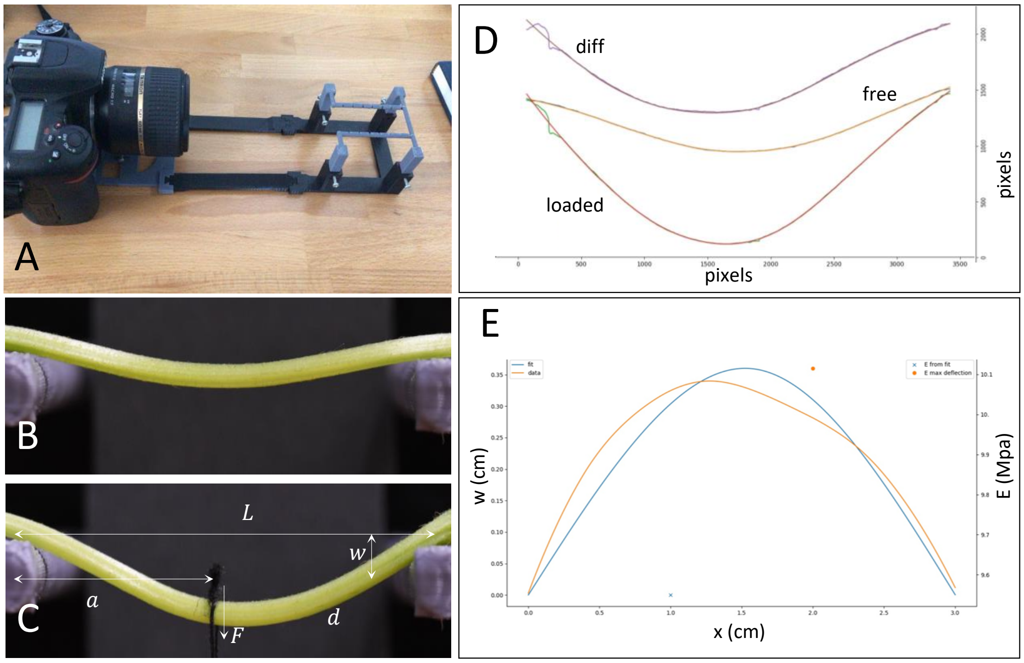


**Figure S13**: **Example of Young's modulus extraction procedure for a stem section after twining experiment using 3-point bending setup. A.** Imaging setup for 3-point bending, custom made with a 3D printer for consistent imaging and stem positioning. **B.** Free stem section (green) and **C.** Stem section for $L - s \in (5 - 10)$ cm, loaded with a weight of 2 g tied to a string (black). Parameters used for Young's modulus extraction shown in figure: $L$ the distance between the supports, $w$ the deflection from the horizontal, $a$ distance of load from left support, $d$ stem diameter, $F$ external force applied with weight. **D.** Extracted skeletons of free and loaded stem sections, and their difference. **F.** Fit of analytic 3-point bending displacement function(blue curve, Eq. S1) to the displacement difference (orange curve), and the extracted Young's moduli from the fit (orange point from max deflection, blue point from fit parameters).

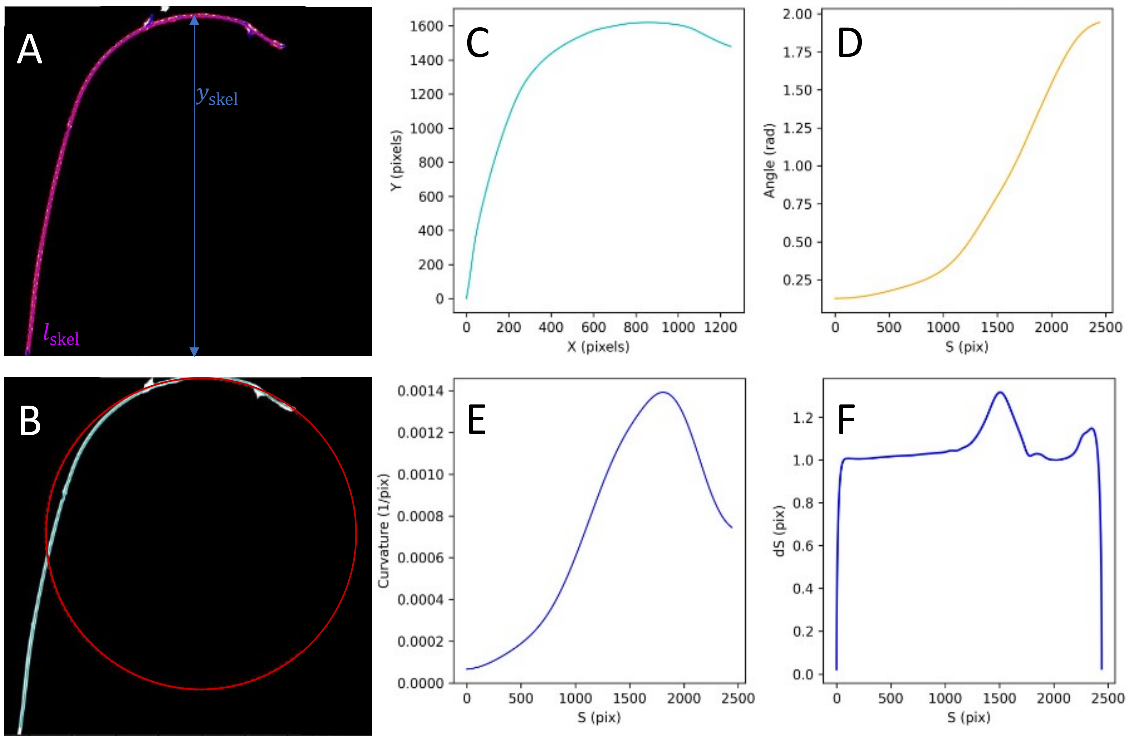


**Figure S14**: **Example of curvature extraction process for stem from side view image during CN.** **A.** Extracted skeleton (magenta) from segmented image.**B.** Circle of radius $R = 1/\max(\kappa)$ overlaid onto skeleton coordinates at the point of maximal curvature. **C.** Extracted skeleton coordinates with origin at the basal part of the stem. **D.** Smoothed angle $\theta$ along the arc length $S$, extracted from skeleton coordinates. **E.** Stem curvature $\kappa$ along the arc length $S$, extracted from the smoothed angle. **F.** Arc length difference $dS$ along the arc length.

**Caption for Movie S1. Twining on a pendulum support.** Before contact, the stem performs free circumnutation and is positioned to naturally encounter the pendulum support (3.1 g). After 1.5 h of contact, stem orientation changes. Twining initiation is defined when the angle of the stem segment contacting the support reaches 45°.

**Caption for Movie S2. Side-view circumnutation snapshot.** Red dots painted along the side of the stem facing the camera continue to face the camera throughout the circumnutating moment, highlighting azimuthal bending, and consistent with the assumption of the model of a fixed material frame.

**Caption for Movie S3. Simulation visualization**. Side (left panel) and top (right panel) views of numerical simulation of organ contacting cylindrical support. Red arrow represents the force vector applied to the cylinder. Absolute curvature difference $\kappa - \kappa_0$ represented by color, values shown in colorbar (right).

**Caption for Movie S4. Motorized experiment.** Top view snapshot of motorized experiment where motor rotation was opposite and approximately equal to the direction of the natural CN rate of the plant. In this case force (and torque) cannot increase, the required deformation is never reached, and the plant never twines, over 12 hours. Once motor rotation is stopped, twining resumes naturally (approximately one hour).
